# Increased Cerebral Serum Amyloid A2 and Parameters of Oxidation in Arylsulfatase B (N-Acetylgalactosamine-4-Sulfatase)-Null Mice

**DOI:** 10.1101/2023.04.03.535377

**Authors:** Sumit Bhattacharyya, Joanne K. Tobacman

## Abstract

**Introduction:** Chondroitin sulfate and chondroitin sulfate proteoglycans have been associated with Alzheimer’s Disease (AD), and the impact of modified chondroitin sulfates is being investigated in several animal and cell-based models of AD. Published reports have shown the role of accumulation of chondroitin 4-sulfate and decline in Arylsulfatase B (ARSB; B-acetylgalactosamine-4-sulfatase) in other pathology, including nerve injury, traumatic brain injury, and spinal cord injury. However, the impact of ARSB deficiency on AD pathobiology has not been reported, although changes in ARSB were associated with AD in two prior reports. The enzyme ARSB removes 4-sulfate groups from the non-reducing end of chondroitin 4-sulfate and dermatan sulfate and is required for their degradation. When ARSB activity declines, these sulfated glycosaminoglycans accumulate, as in the inherited disorder Mucopolysaccharidosis VI.

**Methods:** Reports about chondroitin sulfate, chondroitin sulfate proteoglycans and chondroitin sulfatases in Alzheimer’s Disease were reviewed. Measurements of SAA2, iNOS, lipid peroxidation, chondroitin sulfate proteoglycan 4, and other parameters were performed in cortex and hippocampus from ARSB-null mice and controls by QRT-PCR, ELISA, and other standard assays.

**Results:** SAA2 mRNA expression and protein, CSPG4 mRNA, chondroitin 4-sulfate and i-NOS were increased significantly in ARSB-null mice. Measures of lipid peroxidation and redox state were significantly modified.

**Discussion:** Findings indicate that decline in ARSB leads to changes in expression of parameters associated with AD in the hippocampus and cortex of the ARSB-deficient mouse.

**Conclusions:** Further investigation of the impact of decline in ARSB on the development of AD may provide a new approach to prevent and treat AD.

## INTRODUCTION

The enzyme N-acetylgalactosamine-4-sulfatase (Arylsulfatase B, ARSB) removes 4-sulfate groups from the N-acetylgalactosamine 4-sulfate residues at the non-reducing end of the sulfated glycosaminoglycans chondroitin 4-sulfate (C4S) and dermatan sulfate (DS). It is required for their degradation, and inborn deficiency of ARSB leads to Mucopolysaccharidosis (MPS) VI with accumulation of C4S and DS throughout the body. Disease manifestations include short stature, skeletal abnormalities, reduced life expectancy, corneal opacities, and organ dysfunction. Cognitive decline has been reported [1,2], although not consistently [3]. Decline in ARSB has been detected in bronchial epithelial cells [4] and leukocytes [5] from cystic fibrosis and from malignant cells and tissues of prostate, mammary, colon, and melanoma malignancies [4]. Decline in ARSB was detected in brain tissue from a model of traumatic brain injury [6] and in nerve injury models, including injury of spinal cord [7,8], peripheral nerve [9,10], and optic nerve [11]. These models have accumulation of C4S, and treatment with recombinant ARSB improves functional recovery. ARSB treatment increased neuronal outgrowth following ethanol exposure [12], and affected regrowth of sympathetic nerves following cardiac injury [13]. Decline in ARSB impaired degradation of C4S, and the accrual of C4S impeded regrowth of proper neuronal connections; treatment with rhARSB enabled restoration of the appropriate conduits.

Genetic evidence for the effect of ARSB on development of AD has previously been reported in two published reports. These include a GWAS study of 381 participants in the Alzheimer’s Disease Neuroimaging Initiative in which ARSB was identified as a candidate gene for sporadic AD with a p-value of 6.71 x 10^-6^ [14]. A second report with gene expression profiles in a mouse model of AD showed upregulation of ARSB in surviving neurons in cortical tissue, in contrast to areas with increased Aβ peptide and neuronal loss [15].

For activity, ARSB must undergo post-translational modification, which requires molecular oxygen and the formylglycine generating enzyme (FGE) [16,17]. Investigations indicated that decline in ARSB replicate the effects of hypoxia in human bronchial epithelial and intestinal epithelial cells [18], and that ARSB-mediated changes in chondroitin 4-sulfation affect critical signaling and transcriptional events [4,19], as well as the availability of vital signaling molecules [4]. These findings in different tissues indicate the importance of ARSB in the response to injury and demonstrate how decline in ARSB contributes to diverse pathophysiological mechanisms.

The rationale for these studies was based on several findings, including: 1) the association of decline in ARSB with spinal cord injury, peripheral and optic nerve injury, and traumatic brain injury [6–12]; 2) genetic evidence of association of ARSB with AD [14,15]; 3) previous studies of chondroitin sulfates and chondroitin sulfate proteoglycans in studies of AD [20–26]; and 4) changes in SAA2 expression identified in the hepatic tissue of ARSB-null mice [4], the studies were undertaken to assess whether decline in ARSB and the subsequent increase in C4S in the ARSB-null mouse were associated with parameters related toAD. Based on reports of associations with i-NOS [27,28], glutathione levels [29,30], and lipid peroxidation [31,32], measurements of inducible nitric oxide synthase (iNOS), thiol levels, and lipid peroxidation products, as well as control serum amyloid A2 in cortical and hippocampal tissue of the ARSB-null mouse are reported. Study findings suggest that acquired decline in ARSB, as from impaired oxygenation, may contribute to the pathophysiology of AD.

## METHODS AND MATERIALS

### ARSB null mice

Heterozygous arylsulfatase B deficient mice were obtained (Strain 005598; Jackson Laboratories, Bar Harbor, Maine) and bred, as previously reported [20]. All animal procedures were approved by the Institutional Animal Care and Use Committees (IACUC) of the University of Illinois at Chicago and the Jesse Brown VAMC, Chicago, IL. ARSB defective male mice at 15-17 weeks and age- and gender-matched C57BL/6J control mice were euthanized by carbon dioxide inhalation and cervical dislocation, and the brain was isolated, excised, and promptly frozen at –80°C. Subsequently, hippocampi and cerebral cortex were collected and homogenized for ARSB activity, chondroitin 4-sulfate measurements, and other studies.

### Measurement of ARSB activity

Arylsulfatase B (ARSB) measurements were performed using a fluorometric assay, following a standard protocol with 20 μl of homogenate and 80 μl of assay buffer (0.05 M Na acetate buffer with barium acetate, pH 5.6) with 100 μl of substrate [5mM 4-methylumbelliferyl sulfate (MUS) in assay buffer] in a black microplate, as previously reported [1]. To verify that silencing of ARSB was not associated with generalized reduction of sulfatase activity, galactose-6-sulfatase (GALNS), arylsulfatase A (ARSA), and steroid sulfatase activities were also determined, as previously [4].

### Measurement of chondroitin 4-sulfate

Chondroitin 4-sulfate antibody (C4S; Ly111; Amsbio, Lake Forest, CA) was used to immunoprecipitate C4S from cell lysates of treated and control cells, as previously reported [13]. The Blyscan™ (Biocolor Ltd, Newtownabbey, Northern Ireland) assay was then used to determine the amount of C4S present in the tissues.

### Expression of serum amyloid A2 (SAA2), inducible nitric oxide synthetase (i-NOS) and chondroitin sulfate proteoglycan 4 (CSPG4; neuron-glial antigen 2 or NG2)

SAA2 protein was measured in the hippocampal and cortical tissue and serum of the ARSB null mice and controls using a commercial ELISA (MBS4502344, My BioSource, San Diego, CA) following the instructions. For evaluation of iNOS, SAA2, and CSPG4 gene expression, RT-PCR was performed for mouse SAA2 (NM_011314, mouse iNOS (NM_010927.4), and mouse CSPG4 (NM_00139001.2; NG2) using the primers: SAA2 left: 5’-GGCAGTCCAGGAGGTCTGTA-3’ and right: 5’-GCGAGCCTACACTGACATGA-3’;iNOS left: 5’-CACCTTGGAGTTCACCCAGT-3’ and right: 5’-ACCACTCGTACTTGGGATGC-3’;CSPG4 left: 5’-TACCTGAGCACTGACCCACA-3’ and right: 5’-TTCCCTCTTCCTCCTCTTCC-3’.

### Lipid peroxidation assay

The lipid peroxidation products malonyldialdehyde (MAE) and hydroxyalkenal (HAE) were measured in cerebral cortex of ARSB-null and control mice by a commercial assay (Enzo Life Sciences, Farmingdale, NY), based on the reaction of a chromogenic reagent, N-methyl-2-phenylindole, yielding malondialdehyde (MDA) and 4-hydroxyalkenals (HAE) at 45°C in the presence of methanesulfonic acid. The chromophore was measured at 586 nm.

### Total, inorganic, and protein sulfhydryl determinations

Cellular sulfhydryl in the control and treated samples were determined by a commercial assay (Molecular Probes, Eugene, OR), as previously [19]. Inorganic thiols in the samples reduced a disulfide bond, releasing the active enzyme papain from its inactive S-S form. The activity of the enzyme was then measured using the chromogenic papain substrate, *N*-benzoyl-L-arginine, *p*-nitroanilide (L-BAPNA). The intensity of the developed color was proportionate to the quantity of thiols in each sample. To measure the total (inorganic and protein-associated) thiols, cystamine was added to the reaction, which permitted the detection of poorly accessible thiols on proteins with high pKa values.

### Statistical analysis

Differences between control and ARSB-null mouse tissue samples were determined by unpaired t-tests, two-tailed, with unequal variance and by Mann-Whitney U-tests for non-parametric data for each assay. Six independent samples were analyzed for each assay unless stated otherwise. Box-whisker plots are used to display the data and show the median, minimum, maximum, and quartiles and demonstrate the normality or skewness of the data. Data from controls and from ARSB-null mice separated into two distinct groups, and rank ordering yielded U-values=0 and p-values <0.01. P-values <0.05 are considered statistically significant, and *** indicates p<0.001.

## RESULTS

### Negligible Arylsulfatase B activity and marked increase in chondroitin 4-sulfate in hippocampus and cortex of ARSB null mice

ARSB activity and content of chondroitin 4-sulfate were determined in the cortex and hippocampi of ARSB null mice and controls (**Fig.1A,1B**). As anticipated, ARSB was negligible and C4S was significantly greater in the ARSB null mice than the controls.

**Fig.1.**
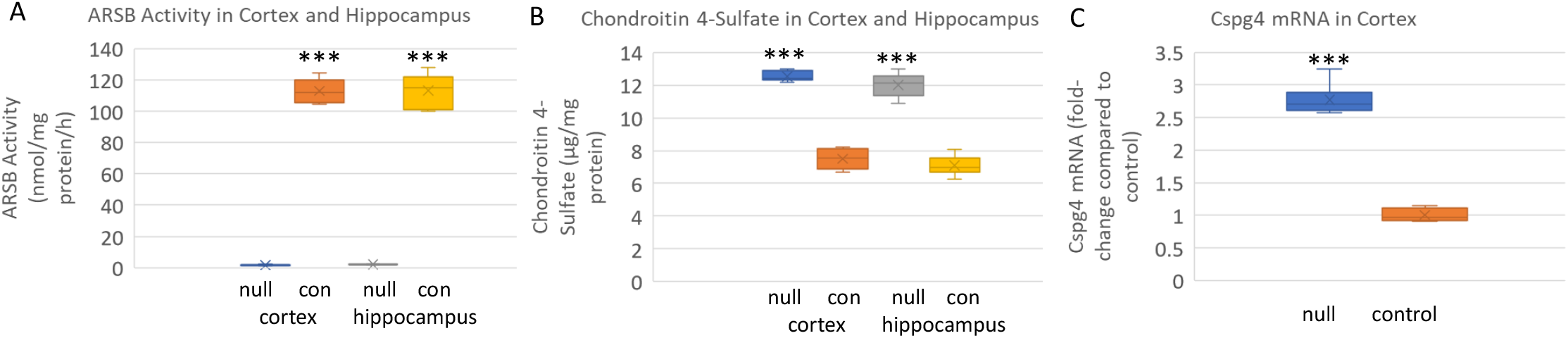
Arylsulfatase B (ARSB) and chondroitin 4-sulfate (C4S) in cortex and hippocampus of ARSB null mice and controls. **A.** ARSB activity was significantly reduced in the brain tissue of the ARSB-null mice, as expected. Level in the control mice was 113.1 ± 10.8 nmol/mg protein/h and 2.1 ± 0.2 nmol/mg protein/h (n=6 in each group) in hippocampal tissue and similar in cortical tissue. **B.** Consistent with the ARSB activity, the C4S in the hippocampus of the ARSB-null mice was 12.0 ± 0.7 μg/g protein and 7.1 ± 0.6 μg/g protein in the control mice, with similar values in the cortical tissue (n=6 in each group). **C.** mRNA expression of chondroitin sulfate proteoglycan 4 (CSPG4) increased to 2.8 times the control level in the cortex of the ARSB-null mice (n=6 in each group). *** for p<0.001 by unpaired t-test, two-tailed, unequal variance; p<0.01 by Mann-Whitney U test for non-parametric data. [con=control; null=ARSB-null]

### Increased expression of chondroitin sulfate proteoglycan 4 in cortex of ARSB null mice

The chondroitin sulfate proteoglycan 4 (CSPG4), also known as NG2 (neuron-glial antigen 2) mRNA expression increased to 2.8 times the baseline level in the brain cortical tissue of the ARSB-null mice, compared to the control (**Fig.1C**).

### Increased SAA2 expression in cortex and hippocampus of ARSB null mice

Investigation of SAA2 gene expression in hippocampus and cortex of male and female ARSB null mice showed increases in SAA protein (**Fig.2A**) and mRNA (**Fig.2B**) concentrations. No difference was apparent between male and female mice. Serum SAA2 levels were similar to values in age-matched C57BL/6J mice (**Fig.2C**).

**Fig.2.**
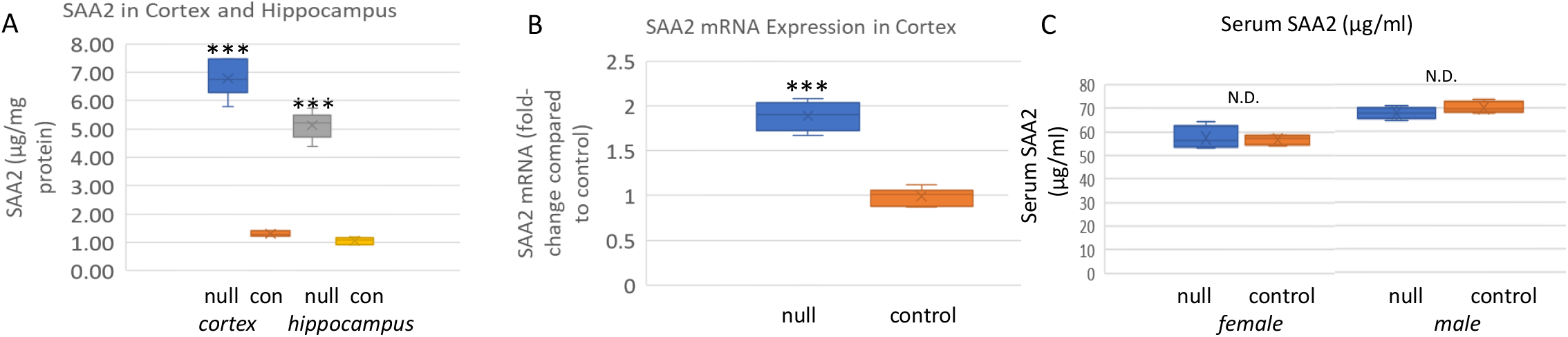
SAA in cortex, hippocampus, and serum of ARSB null mice and controls. **A.** SAA was measured by ELISA in the cortex and hippocampus of control and ARSB null mice. In cortex, control value was 1.30 ± 0.09 μg/mg protein compared to 6.78 ± 0.63 μg/mg protein in the ARSB-null mouse cortex. Values in hippocampus were similar (1.06 ± 0.12 vs. 5.13 ± 0.47 μg/mg protein; n=6 in all groups). P-values are <0.001 for all comparisons (unpaired t-test, two-tailed). **B.** QRT-PCR data indicate 1.89-times higher expression in the ARSB-null mice than the control mice (n=6; p<0.001, unpaired t-test, two-tailed). Values were similar for male and female mice. **C**. Serum levels of SAA were similar in control and ARSB-null mice, and higher in the male mice than the female mice (n=4 in each group). [con=control; null=ARSB-null]

### Increased expression of inducible nitric oxide synthase in cortex of ARSB null mice

i-NOS was measured in hippocampal and cortical tissue of the ARSB null and control mice (**Fig.3A**). Results indicated marked increase in iNOS expression in the ARSB null mice

**Fig.3.**
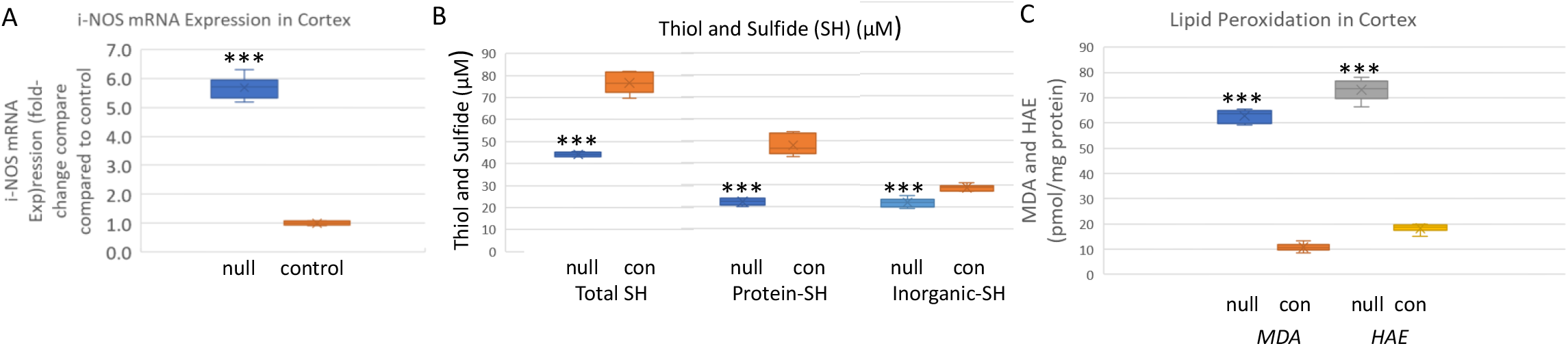
Inducible nitric oxide synthase, total thiols, and products of lipid peroxidation are increased in the cortex of ARSB-null mice. **A.** Inducible nitric oxide synthase (i-NOS) expression was markedly increased in the ARSB-null mice (5.68 ± 0.39 vs. 1.04 ± 0.08, n=6; p<0.001, unpaired t-test, two-tailed, unequal variance). **B.** Total thiols, including protein-thiol groups and inorganic SH groups were lower in the ARSB-null mice than the controls (71.6 ± 4.9 vs. 44.1 ± 1.0; n=6, p<0.001), consistent with lower reducing capacity in the ARSB-null mice. **C.** The lipid peroxidation products, MDA and HAE, were measured in the cortical tissue of the ARSB-null and control mice. In the ARSB-null mouse tissue, levels are several-fold higher (n=6, p<0.001). [con=control; HAE=4-hydroxyalkenals; MDA=malondialdehyde; null=ARSB-null; SH=sulfhydryl]

### Decline in sulfhydryls in cortex of ARSB null mice

Decline in ARSB was previously associated with marked changes in the overall redox status of human bronchial and colonic epithelial cells. Findings indicated that when ARSB was silenced by siRNA, the redox capacity of the cells was markedly affected and levels of reduced glutathione and of thiols were significantly reduced [19]. In cortical tissue from the ARSB null mice, the levels of total, organic, and inorganic sulfide and thiol, markers of overall cellular redox activity, were markedly reduced (**Fig.3B**).

### Increase in lipid oxidation products in cortex of ARSB null mice

Lipid peroxidation is a well-established indicator of oxidative stress in cells and tissues. Polyunsaturated fatty acid peroxides generate malondialdehyde (MDA) and 4-hydroxyalkenals (HAE) upon decomposition, and measurements of MDA and HAE indicate extent of lipid peroxidation. In association with the marked change in redox state, the levels of HAE and MDA were significantly increased in cortical tissue of the ARSB null mice (**Fig.3C**). These increases reflect enhanced lipid peroxidation in the absence of ARSB activity.

## DISCUSSION

Exploration of the etiology of Alzheimer’s dementia is ongoing, engaging laboratories in thousands of basic, translational, and clinical investigations. To this prodigious research endeavor, this short communication contributes new, basic information linking decline in ARSB with increased hippocampal and cortical SAA2 in the mouse brain. Corresponding to these data are increases in expression of i-NOS and lipid peroxidation products and decline in abundance of sulfhydryls, consistent with the overall finding of heightened oxidative process in the AD brain [27–32]. The study findings are consistent with previous reports of GWAS data linking ARSB and increased AD [14] and of relative increase in ARSB in neurons lacking amyloid in AD brains [15]. Study data are also consistent with previous findings of increase in chondroitin sulfate proteoglycans, specifically CSPG4 [4], neurocan [12], and versican [19], and chondroitin 4-sulfate [4,12,18,19], when ARSB was silenced in human cells.

Interestingly, although there are significant increases in the cerebral SAA2 in the ARSB-null mice, serum SAA2 is not changed and the hepatic expression of SAA2 (NM_011314) and SAA1 (NM_009117) was markedly decreased in a gene expression array [33]. This corresponds to a report of lower Aβ levels in liver samples from AD patients compared to healthy controls [34]. The data are consistent with a role of the liver in the clearance of Aβ peptides and with a complex, integrated, systemic response to decline in ARSB and increase in amyloid.

Several reports demonstrate a critical impact of decline in ARSB and the associated increase in chondroitin 4-sulfate on nerve injury [7–12]. Similar buildup of C4S may contribute to CNS dysfunction due to impaired neuronal signaling. Dependence of ARSB on oxygen for function, due to post-translational modification of ARSB requiring oxygen and the formylglycine generating enzyme [16], and replication of the effects of hypoxia by decline in ARSB [18], suggest that impaired circulation and oxygen delivery can produce the effects of decline in ARSB.

In contrast to MPS VI, in which ARSB is defective, inherited deficiencies of some other sulfatases are characterized by dementia. These disorders include MPS III (Sanfilippo Disease), in which decline in heparan N-sulfatase or N-acetylglucosamino-6-sulfatase affects accumulation of heparan sulfate, and metachromatic leukodystrophy (MLD), in which decline in Arylsulfatase A leads to accumulation of the sulfatides cerebroside sulfate and 3-O-sulfo-galactosylceramide. Congenital deficiency of ARSB in MPS VI is not associated with dementia, and the limited life expectancy of individuals with MPS VI prevents assessment of whether dementia develops in later years. A feline model of ARSB deficiency detected abnormal lysosomal storage in some cortical neurons and glia, which resembled findings in other lysosomal storage disorders [35].

The impact of proteoglycans in AD has been recognized for decades, with much of the early attention focused on heparan sulfate [3,36,37]. Subsequently, investigations have reported about the accumulation of chondroitin sulfates (CS) and chondroitin sulfate proteoglycans (CSPGs) in AD tissues, with over 150 publications. CSPGs were detected in senile plaques and neurofibrillary tangles [20–22], and the degree of sulfation of glycosaminoglycans was identified as critical for the interaction with tau protein [23]. CSPGs are present in the CNS perineuronal nets which regulate plasticity, and C4S was observed to be more inhibitory than C6S to axon growth and regeneration [24,25]. Analysis of chondroitin disaccharides may lead to more precise information about the extent and sites of chondroitin sulfation in AD [26].

Treatment approaches focused on reducing the effects of CS/CSPG have included chondroitinase ABC [38], antibody to aggrecan, a CSPG [39], and inhibition of Aβ-CS interaction by mono- and di-saccharides derived from CS [40]. The potential benefit of low molecular weight chondroitin sulfate (LMWCS) oligosaccharides and modified LMWCS has also been studied [41–44]. Treatments have included selenium-modified CS [45[, lithium-modified CS [46], and gold-modified CS [47]. This approach is based on competition and inhibition of CS/CSPG function by the oligosaccharides. The oligosaccharides are derived from shark cartilage which is composed predominantly of chondroitin 6-sulfate. This approach is consistent with a focus on inhibition of effects of C4S by C6S derivatives.

Recombinant human ARSB is safely used for replacement of defective ARSB in MPS VI and may have a role in AD. Further examination of the role of ARSB in the etiology of AD may lead to new approaches to effective intervention and to clarification of the underlying pathophysiology.

## ACKNOWLEDGMENTS

The authors acknowledge the contributions of Leo Feferman, M.D. to some of the studies in this report. Also, the authors acknowledge use of facilities of the JBVAMC. The content is solely the responsibility of the authors and does not necessarily represent the official views of the NIH or of the VA.

## DATA AVAILABILITY

Data are available by communication with JKT or SB.

## REFERENCES

1. Ebbink BJ, Brands MM, van den Hout JM, Lequin MH, Coebergh van den Braak RR, van de Weitgraven RL, Plug I, Aarsen FK, van der Ploeg AT. (2016) Long-term cognitive follow-up in children treated for Maroteaux-Lamy syndrome. J Inherit Metab Dis 39(2), 285–292.

2. Kılıç M, Dursun A, Coşkun T, Tokatlı A, Özgül RK, Yücel-Yılmaz D, Karaca M, Doğru D, Alehan D, Kadayifçilar S, Genç A, Turan-Dizdar H, Gönüldaş B, Savcı S, Sağlam M, Aksoy C, Arslan U, Sivri HS (2017) Genotypic-phenotypic features and enzyme replacement therapy outcome in patients with mucopolysaccharidosis VI from Turkey. Am J Med Genet A 173(11), 2954–2967.

3. Gomes DF, Gallo LG, Leite BF, Silva RB, da Silva EN (2019). Clinical effectiveness of enzyme replacement therapy with galsulfase in mucopolysaccharidosis type VI treatment: Systematic review. J Inherit Metab Dis 42, 66–76.

4. Tobacman JK, Bhattacharyya S. (2022) Profound impact of decline in N-acetylgalactosamine-4-sulfatase (Arylsulfatase B) on molecular pathophysiology and human diseases. Int J Mol Sci. 23(21), 13146.

5. Sharma G, Burke J, Bhattacharyya S, Sharma N, Katyal S, Park RL, Tobacman J (2013) Reduced arylsulfatase B activity in leukocytes from cystic fibrosis patients. Pediatr Pulmonol 48(3), 236–244.

6. Bhattacharyya S, Zhang X, Feferman L, Johnson D, Tortella FC, Guizzetti M, Tobacman JK (2015) Decline in arylsulfatase B and Increase in chondroitin 4-sulfotransferase combine to increase chondroitin 4-sulfate in traumatic brain injury. J Neurochem 134(4), 728–739.

7. Yoo M, Khaled M, Gibbs KM, Kim J, Kowalewski B, Dierks T, Schachner M. (2013) Arylsulfatase B improves locomotor function after mouse spinal cord injury. PloS One 8(3), e57415.

8. Park HH, Kim YM, Anh Hong LT, Kim HS, Kim SH, Jin X, Hwang DH, Kwon MJ, Song SC, Kim BG (2022) Dual-functional hydrogel system for spinal cord regeneration with sustained release of arylsulfatase B alleviates fibrotic microenvironment and promotes axonal regeneration. Biomaterials 284,121526.

9. Pearson CS, Mencio CP, Barber AC, Martin KR, Geller HM (2018) Identification of a critical sulfation in chondroitin that inhibits axonal regeneration. Elife 7, e37139.

10. Wang H, Katagiri Y, McCann TE, Unsworth E, Goldsmith P, Yu ZX, Tan F, Santiago L, Mills EM, Wang Y, Symes AJ, Geller HM (2008) Chondroitin-4-sulfation negatively regulates axonal guidance and growth. J Cell Sci 121(Pt 18), 3083–3091.

11. Pearson CS, Solano AG, Tilve SM, Mencio CP, Martin KR, Geller HM (2020) Spatiotemporal distribution of chondroitin sulfate proteoglycans after optic nerve injury in rodents. Exp Eye Res 190, 107859.

12. Zhang X, Bhattacharyya S, Kusumo H, Goodlett CR, Tobacman JK, Guizzetti M (2014) Arylsulfatase B modulates neurite outgrowth via astrocyte chondroitin-4-sulfate: dysregulation by ethanol. Glia 62(2), 259–271.

13. Blake MR, Parrish DC, Staffenson MA, Sueda S, Woodward WR, Habecker BA (2022) Chondroitin sulfate proteoglycan 4,6 sulfation regulates sympathetic nerve regeneration after myocardial infarction. Elife 23, 11:e78387.

14. Potkin SG, Guffanti G, Lakatos A, Turner JA, Kruggel F, Fallon JH, Saykin AJ, Orro A, Lupoli S, Salvi E, Weiner M, Macciardi F (2009) Alzheimer’s Disease Neuroimaging Initiative. Hippocampal atrophy as a quantitative trait in a genome-wide association study identifying novel susceptibility genes for Alzheimer’s disease. PLoS One 4(8), e6501.

15. Li Y, Xu C, Schubert D (1999) The up-regulation of endosomal-lysosomal components in amyloid beta-resistant cells. J Neurochem 73(4), 1477–1482.

16. Roeser D, Preusser-Kunze A, Schmidt B, Gasow K, Wittmann JG, Dierks T, von Figura K, Rudolph MG (2006) A general binding mechanism for all human sulfatases by the formylglycine-generating enzyme. Proc Natl Acad Sci USA 103(1), 81–86.

17. Roeser D, Schmidt B, Preusser-Kunze A, Rudolph MG (2007) Probing the oxygen-binding site of the human formylglycine-generating enzyme using halide ions. Acta Crystallogr D Biol Crystallogr 63(Pt 5), 621–627.

18. Bhattacharyya S, Tobacman JK (2012) Hypoxia reduces arylsulfatase B activity and silencing arylsulfatase B replicates and mediates the effects of hypoxia. PLoS One 7(3), e33250.

19. Bhattacharyya S, Feferman L, Tobacman JK (2014) Arylsulfatase B regulates versican expression by galectin-3 and AP-1 mediated transcriptional effects. Oncogene 33(47), 5467–5476.

20. DeWitt DA, Silver J, Canning DR, Perry G (1993) Chondroitin sulfate proteoglycans are associated with the lesions of Alzheimer’s disease. Exp Neurol 121(2), 149–152.

21. Canning DR, McKeon RJ, DeWitt DA, Perry G, Wujek JR, Frederickson RC, Silver J (1993) beta-Amyloid of Alzheimer’s disease induces reactive gliosis that inhibits axonal outgrowth. Exp Neurol 124(2), 289–298.

22. DeWitt DA, Richey PL, Praprotnik D, Silver J, Perry G (1994) Chondroitin sulfate proteoglycans are a common component of neuronal inclusions and astrocytic reaction in neurodegenerative diseases. Brain Res 656(1), 205–209.

23. Hasegawa M, Crowther RA, Jakes R, Goedert M (1997) Alzheimer-like changes in microtubule-associated protein Tau induced by sulfated glycosaminoglycans. Inhibition of microtubule binding, stimulation of phosphorylation, and filament assembly depend on the degree of sulfation. J Biol Chem 272(52), 33118–24.

24. Galtrey CM, Fawcett JW (2007) The role of chondroitin sulfate proteoglycans in regeneration and plasticity in the central nervous system. Brain Res Rev 54(1), 1–18.

25. Foscarin S, Raha-Chowdhury R, Fawcett JW, Kwok JCF (2017) Brain ageing changes proteoglycan sulfation, rendering perineuronal nets more inhibitory. Aging (Albany NY) 9(6), 1607–1622.

26. Logsdon AF, Francis KL, Richardson NE, Hu SJ, Faber CL, Phan BA, Nguyen V, Setthavongsack N, Banks WA, Woltjer RL, Keene CD, Latimer CS, Schwartz MW, Scarlett JM, Alonge KM (2022) Decoding perineuronal net glycan sulfation patterns in the Alzheimer’s disease brain. Alzheimers Dement 18(5), 942–954.

27. Dorheim MA, Tracey WR, Pollock JS, Grammas P (1994) Nitric oxide synthase activity is elevated in brain microvessels in Alzheimer’ s disease. Biochem Biophys Res Commun 205, 659–665.

28. Asiimwe N, Yeo SG, Kim M-K, Jung J, Jeong NY (2016) Nitric Oxide: Exploring the contextual link with Alzheimer’s Disease. Oxid Med Cell Longev 2016, 7205747.

29. Saharan S, Mandal PK (2014) The emerging role of glutathione in Alzheimer’s disease. J Alzheimers Dis 40(3), 519–529.

30. McBean GJ, Aslan M, Griffiths HR, Torrão RC (2015) Thiol redox homeostasis in neurodegenerative disease. Redox Biol 5, 186–194.

31. Montine TJ, Neely MD, Quinn JF, Beal MF, Markesbery WR, Roberts LJ, Morrow JD. (2002) Lipid peroxidation in aging brain and Alzheimer’s disease. Free Radic Biol Med. 33(5), 620–626.

32. Miyazawa T (2021) Lipid hydroperoxides in nutrition, health, and diseases. Proc Jpn Acad Ser B Phys Biol Sci 97(4), 161–196.

33. NCBI.NLM.NIH.gov/GEO DataSets/Arylsulfatase B/GSE77689

34. Roher AE, Esh CL, Kokjohn TA, Castaño EM, Van Vickle GD, Kalback WM, Patton RL, Luehrs DC, Daugs ID, Kuo YM, Emmerling MR, Soares H, Quinn JF, Kaye J, Connor DJ, Silverberg NB, Adler CH, Seward JD, Beach TG, Sabbagh MN (2009) Amyloid beta peptides in human plasma and tissues and their significance for Alzheimer’s disease. Alzheimers Dement 5(1), 18–29.

35. Walkley SU, Thrall MA, Haskins ME, Mitchell TW, Wenger DA, Brown DE, Dial S, Seim H (2005) Abnormal neuronal metabolism and storage in mucopolysaccharidosis type VI (Maroteaux-Lamy) disease. Neuropathol Appl Neurobiol 31(5), 536–544.

36. Snow AD, Wight TN (1989) Proteoglycans in the pathogenesis of Alzheimer’s disease and other amyloidosis. Neurobiol Aging 10, 481–497.

37. Snow AD, Cummings JA and Lake T (2021) The unifying hypothesis of Alzheimers Disease: Heparan sulfate proteoglycans/glycosaminoglycans are key as first hypothesized over 30 years ago. Front Aging Neurosci 13, 710683.

38. Howell MD, Bailey LA, Cozart MA, Gannon BM, Gottschall PE (2015) Hippocampal administration of chondroitinase ABC increases plaque-adjacent synaptic marker and diminishes amyloid burden in aged APPswe/PS1dE9 mice. Acta Neuropathol Commun 3, 54.

39. Yang S, Hilton S, Alves JN, Saksida LM, Bussey T, Matthews RT, Kitagawa H, Spillantini MG, Kwok JCF, Fawcett JW (2017) Antibody recognizing 4-sulfated chondroitin sulfate proteoglycans restores memory in taupathy-induced neurodegeneration. Neurobiol Aging 59, 197–209.

40. Fraser PE, Darabie AA, McLaurin JA (2001) Amyloid-beta interactions with chondroitin sulfate-derived monosaccharides and disaccharides. implications for drug development. J Biol Chem 276(9), 6412–9.

41. Zhang Q, Li J, Liu C, Song C, Li P, Yin F, Xiao Y, Li J, Jiang W, Zong A, Zhang X, Wang F (2015) Protective effects of low molecular weight chondroitin sulfate on amyloid beta (Aβ)-induced damage in vitro and in vivo. Neuroscience 305, 169–182.

42. Zhang Q, Na Z, Cheng Y, Wang F (2018) Low-molecular-weight chondroitin sulfate attenuated injury by inhibiting oxidative stress in amyloid β-treated SH-SY5Y cells. Neuroreport 29(14),1174–1179.

43. Zhao N, Meng J, Jiang W, Xu W, Liu C, Wang F (2021) Study on the relationships between molecular weights of chondroitin sulfate oligosaccharides and Aβ-induced oxidative stress and the related mechanisms. Glycobiology 31(4), 492–507.

44. Zhao N, Wu L, Zhang X, Jiang W, Wang F (2021) Low molecular weight chondroitin sulfate ameliorates pathological changes in 5XFAD mice by improving various functions in the brain. Neuropharmacology 199, 108796.

45. Ji D, Wu X, Li D, Liu P, Zhang S, Gao D, Gao F, Zhang M, Xiao Y (2020) Protective effects of chondroitin sulphate nano-selenium on a mouse model of Alzheimer’s disease. Int J Biol Macromol 154, 233–245.

46. Gao D, Li P, Gao F, Feng Y, Li X, Li D, Li Y, Xiao Y (2022) Preparation and multitarget anti-AD activity study of chondroitin sulfate lithium in AD mice induced by combination of D-Gal/AlCl3. Oxid Med Cell Longev 12, 9466166.

47. Feng Y, Li X, Ji D, Tian J, Peng Q, Shen Y, Xiao Y (2023) Functionalised penetrating peptide-chondroitin sulphate-gold nanoparticles: Synthesis, characterization, and applications as an anti-Alzheimer’s disease drug. Int J Biol Macromol 230, 123125.

